# xMD-miRNA-seq to generate near in vivo miRNA expression estimates in colon epithelial cells

**DOI:** 10.1101/333658

**Authors:** Avi Z. Rosenberg, Carrie Wright, Karen Fox-Talbot, Anandita Rajpurohit, Courtney Williams, Corey Porter, Olga Kovbasnjuk, Matthew N. McCall, Joo Heon Shin, Marc K. Halushka

## Abstract

Accurate, RNA-seq based, microRNA (miRNA) expression estimates from primary cells have recently been described. However, this *in vitro* data is mainly obtained from cell culture, which is known to alter cell maturity/differentiation status, significantly changing miRNA levels. What is needed is a robust method to obtain *in vivo* miRNA expression values directly from cells. We introduce expression microdissection miRNA small RNA sequencing (xMD-miRNA-seq), a method to isolate cells directly from formalin fixed paraffin-embedded (FFPE) tissues. xMD-miRNA-seq is a low-cost, high-throughput, immunohistochemistry-based method to capture any cell type of interest. As a proof-of-concept, we isolated colon epithelial cells from two specimens and performed low-input small RNA-seq. We generated up to 600,000 miRNA reads from the samples. Isolated epithelial cells, had abundant epithelial-enriched miRNA expression (miR-192; miR-194; miR-200b; miR-200c; miR-215; miR-375) and overall similar miRNA expression patterns to other epithelial cell populations (colonic enteroids and flow-isolated colon epithelium). xMD-derived epithelial cells were generally not contaminated by other adjacent cells of the colon as noted by t-SNE analysis. xMD-miRNA-seq allows for simple, economical, and efficient identification of cell-specific miRNA expression estimates. Further development will enhance rapid identification of cell-specific miRNA expression estimates in health and disease for nearly any cell type using archival FFPE material.

## Background

MicroRNAs (miRNAs) are small, regulatory RNA elements with critical control of protein expression. Most miRNAs are well-conserved between species with expression patterns that vary during development and disease.^1,2^ Three cell-focused manuscripts recently described miRNA expression at the cell level, rather than at the tissue level.^3-5^ This cell-specific expression knowledge is critical to understand the important mechanistic activity of miRNAs as they relate to disease.^6,7^ To date, the majority of our cell-specific expression miRNA knowledge comes from primary cell culture. However, this source has significant limitations.

*In vitro* cell culture causes considerable phenotypic changes to a cell. Typically, high serum levels drive cells to proliferate rapidly rather than maintaining a quiescent, mature state.^8^ Without co-cultures, cells also lose important cell-cell interactions and alter their phenotype. Therefore, it is well-established that cultured cells are good, but not ideal surrogates for *in vivo* expression.^9^ This was nicely demonstrated for miRNAs in a study that compared primary endothelial cells directly harvested from umbilical cords to endothelial cells cultured for 3 passages. miR-126, a highly-expressed, mature endothelial cell miRNA, was over 2 fold less abundant at passage 3 versus passage 0. Conversely, several proliferation-related miRNAs of the miR-17-92 cluster were upregulated 3-6 fold over the same time course.^10^ These cell culture-mediated aberrations in relative miRNA expression levels can greatly impact disease-related studies.

There has been a burgeoning interest in deconvoluting tissues into their cellular components for the purpose of better analyzing disease expression datasets and extracting meaningful disease driven cellular changes.^11^ Cellular composition of tissues is highly variable between samples, even when all samples share the same phenotype.^12^ A robust way to deconvolute a tissue is to utilize an expression matrix of each composite cell type to computationally separate the tissue into each individual cell type.^13,14^ For that purpose, expression estimates must closely hew to *in vivo* data. We have noted that often cell-culture based expression estimates fail in this capacity. For example, the reads per million miRNA reads (RPM) value of miR-200c, an epithelial cell specific miRNA, was ~60,000 RPM in multiple human bladder samples. In the bladder, the only native epithelial cell type, representing ~20-80% of a bladder biopsy, is the urothelial cell. However, urothelial cells grown in culture demonstrate a miR-200c value of only 5,000 RPM. It is difficult to reconcile this difference other than to acknowledge that this miRNA, associated with a mature cell phenotype, is greatly reduced in a cell-culture sample.^15^

To overcome this problem, there is a need for methods to capture *in vivo* cell expression miRNA estimates in a robust and cost-effective manner. Excellent methods to obtain cells directly from tissues exist, but each has limitations. Laser-capture microdissection is expensive, tedious, and can only capture sufficient numbers of a particular cell type if they form large structures (ex. glands); otherwise the background contamination of neighboring cells is rate-limiting.^16^ Flow capture and magnetic bead separation are useful for tissues that easily dissociate (ex. blood, but not heart), but these methods are also limited by the widely variable miRNA expression that can occur as a result of methodologic manipulation.^4,15,17^ Single-cell sequencing has great promise, however current methodologies are limited for miRNAs due to cost, and depth of sequencing per cell.^18^

We have previously utilized expression microdissection (xMD) to isolate prostate stroma and epithelium and assay miRNA by droplet digital PCR (ddPCR).^19^ That study led us to hypothesize we could obtain adequate RNA yields for a global survey of miRNA levels by small RNA-sequencing (RNA-seq). We now introduce xMD-miRNA-seq, a method to obtain nearly *in vivo* miRNA expression estimates from any cell type directly from formalin-fixed paraffin-embedded (FFPE) tissues by utilizing expression microdissection.^20^ We demonstrate this technique as an efficient, robust and cost-effective method for generating deep sequencing data to provide accurate miRNA expression data from cells.

## Results

### Successful isolation of epithelial cells by xMD

We performed xMD on six slides from two normal colon samples, performing immunohistochemistry (IHC) for the epithelial-specific cytokeratin marker AE1/AE3. The xMD method yielded a highly enriched population of colonic epithelial cells (Figure 1), with an overall capture rate of approximately 20% of the stained cells onto the ethylene vinyl acetate (EVA) membrane. Total RNA extraction and purification was performed on the cells annealed to the EVA membranes yielding 266 (13.3 ng/μl) and 374 (18.7 ng/μl) ng of total RNA respectively for the two samples. These RNA samples were split into two technical replicates for small RNA-seq.

**Figure 1:**
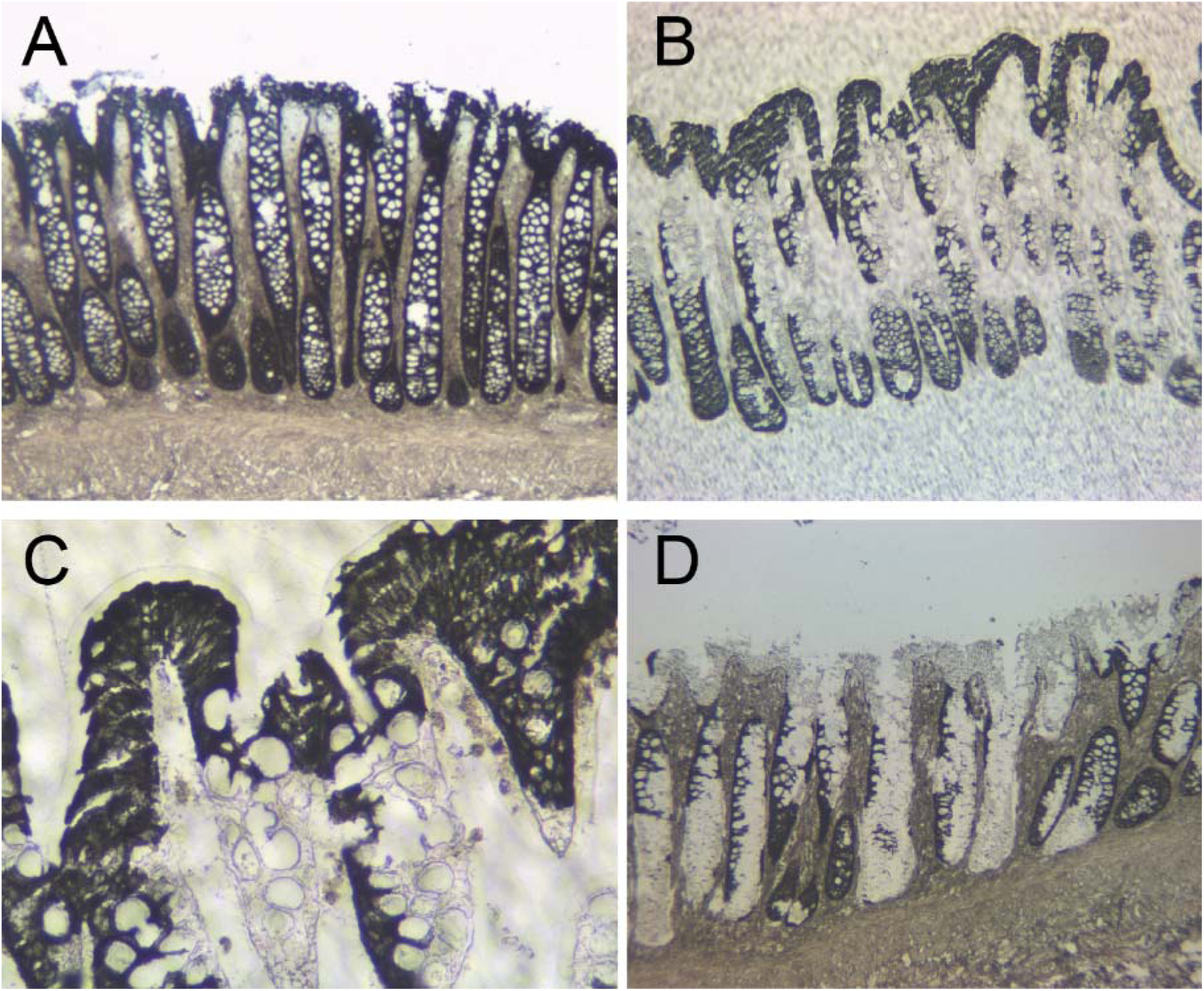
Isolation of colonic epithelial cells using xMD. A) A slide undergoes IHC for the cytokeratin AE1/AE3, a common antibody for epithelial cells. This slide is not coverslipped or counterstained. B) After xMD, the pigmented cells are selectively transferred onto an EVA membrane. C) High-power view of the EVA membrane demonstrating the tight isolation of the pigmented epithelial cells. D) The original slide, after xMD, shows an incomplete transfer of epithelial cells, but no transfer of lamina propria or stromal areas (Images in A, B & D are 100x original magnification while image C is 400x original magnification).

### xMD-miRNA-seq results

Small RNA-seq libraries were generated from 50 ng of total RNA in duplicate for both samples using the BIOO Scientific NEXTflex small RNA library preparation kit V3 for Illumina and the Perkin Elmer Sciclone G3. Final library concentrations measured by a ThermoFisher Qubit Fluorometer 2.0 were 0.26 ng/μl and 0.24 ng/μl for the replicates of the first sample and 0.22 ng/μl and 0.25 ng/μl for the replicates of the second sample. Given these low concentrations, the libraries were sequenced with high depth on an Illumina HiSeq 3000. Adapters and random base linkers were removed with Cutadapt version 1.14 and trimmed FASTQ files were processed using both miRge and a miRBase v21 miRNA library and miRge 2.0 with a MirGeneDB library of miRNAs.^21-24^ Using the more restrictive MirGeneDB dataset, we identified between 260 and 321 miRNAs for each sample. Setting the minimum threshold to 10 RPM for a given miRNA, the range was 236-285 miRNAs per sample, with 203 miRNAs consistently above 10 RPM. The number of miRNA reads varied between 172,678 and 616,921 (Table 1). As a percent of all reads, these yields were low (0.54-0.65% of total reads); requiring deep sequencing of each sample with the current method. Within the sequencing reads, both rRNA sequences and tRNA halves and fragments were abundant accounting for approximately 30-45% and 10-15% of reads respectively.

**Table 1:** xMD-miRNA-seq read composition by RNA type.

| xMD Sample | Total Input Reads | Filtered miRNA Reads | Unique miRGeneDB miRNAs | rRNA Reads | Other ncRNA Reads | mRNA Reads | Remaining Reads | % miRNAs |
| --- | --- | --- | --- | --- | --- | --- | --- | --- |
| xMD-Epi-1 | 78,393,406 | 430,919 | 281 | 25,217,820 | 3,626,007 | 877,321 | 24,759,053 | 0.55% |
| xMD-Epi-2 | 31,908,081 | 172,678 | 260 | 10,676,670 | 1,501,036 | 361,505 | 11,080,747 | 0.54% |
| xMD-Epi-3 | 39,826,926 | 248,093 | 264 | 17,000,621 | 2,077,002 | 642,129 | 13,849,697 | 0.62% |
| xMD-Epi-4 | 95,396,126 | 616,921 | 321 | 40,012,705 | 5,114,006 | 1,535,607 | 32,500,021 | 0.65% |

A number of technical factors were investigated to better understand the variability in xMD-miRNA-Seq. Technical duplicates of the four samples showed similar expression patterns with Pearson correlation r values of log_2_ adjusted data of 0.83 and 0.88 respectively (Supplementary Figure 1A). The length of xMD-derived miRNAs were similar to endothelial (SRR5127228) and fibroblast (SRR5127231) primary cells grown in culture although with more shorter reads; whereas epithelial cells that were flow-collected (SRR5127219) had a much higher abundance of shorter miRNA reads (Supplementary Figure 1B). We then assessed which miRNAs varied the most between technical replicates. Using only miRNAs with a minimum average RPM >1000 (N=90) we noted only sixteen miRNAs had >4 fold change difference. Not surprisingly the fold change differences were highest in samples with the lowest RPM values, with two miRNAs, miR-532-5p and miR-425-5p being the most variable (Supplementary Figure 1C). Finally, we investigated the coefficient of variation across these 90 miRNAs finding a trend for a higher coefficient of variation for the lower abundance miRNAs. However, no miRNAs were particular outliers (Supplementary Figure 1D). Due to the lack of significant outliers, no structural features such as %GC or positional nucleotide usage that may vary and affect miRNA levels were noted.^25^

### A highly enriched epithelial signal

Each sample was investigated for its most abundant miRNAs (Table 2). On average, the most abundant miRNA in each sample was the combined let-7a-5p/let-7c-5p miRNA. Following those were five well-known epithelial specific/enriched miRNAs (miR-215-5p/192-5p; miR-375; miR-200c-3p; miR-200b-3p; miR-194) in the top twenty most abundant miRNAs. Five of these miRNAs are generic epithelial markers, but miR-375 is specifically enriched in colonic epithelium in addition to being the most highly expressed miRNA in pancreatic islet cells.^26^ Two miRNAs likely represent different sources of contamination. miR-143-3p, which is a highly-abundant mesenchyme-enriched miRNA, has generally been shown to have low^4^ to absent epithelial expression.^27^ We surmise this is a contaminant resulting from capturing some fibroblasts or smooth muscle cells on the EVA membranes. miR-9-5p is a neural enriched miRNA with trivial expression in colon.^4^ This signal may have been the result of differences in NEXTflex and Illumina TruSeq library preparation kits that favor higher counting of this miRNA in the NEXTflex system or contamination from library preparation with brain samples that were concurrently sequenced, as has been described.^28^ The remaining miRNAs are well-known and ubiquitously expressed across many cell types.

**Table 2:** Top twenty most abundant miRNAs by RPM value for four replicates of xMD-miRNA-seq derived epithelial cells. Five miRNAs are epithelial specific (+), while two (-) reflect contamination. Of these, xMD-Epi-1 and xMD-Epi-2 are technical replicates as are xMD-Epi-3 and xMD-Epi-4. miR-215-5p/192-5p and miR-27a-3p/27b-3p are shown together due to the similarities of their sequences and conservative approach of miRge to assign isomiR reads.

| miRNA | xMD-Epi-1 | xMD-Epi-2 | xMD-Epi-3 | xMD-Epi-4 | Epithelial Enriched |
| --- | --- | --- | --- | --- | --- |
| let-7a-5p/7c-5p | 89,237 | 81,591 | 60,586 | 63,122 |  |
| miR-215-5p/192-5p | 66,166 | 45,680 | 43,367 | 62,781 | + |
| miR-26a-5p | 45,777 | 65,503 | 50,852 | 53,801 |  |
| miR-92a-3p | 40,003 | 52,439 | 47,668 | 42,454 |  |
| miR-375 | 40,103 | 40,376 | 40,231 | 48,419 | + |
| miR-200c-3p | 36,701 | 27,462 | 44,894 | 37,984 | + |
| miR-191-5p | 33,025 | 31,712 | 36,575 | 39,449 |  |
| let-7f-5p | 35,181 | 31,174 | 27,594 | 29,837 |  |
| let-7b-5p | 27,260 | 17,895 | 38,647 | 32,806 |  |
| miR-200b-3p | 30,412 | 25,018 | 29,348 | 28,527 | + |
| miR-21-5p | 26,982 | 30,566 | 26,458 | 25,622 |  |
| miR-143-3p | 34,584 | 29,106 | 20,146 | 24,493 | - |
| miR-9-5p | 16,061 | 30,872 | 24,438 | 27,976 | - |
| let-7g-5p | 24,172 | 17,796 | 29,437 | 24,583 |  |
| miR-328-3p | 46,900 | 37,758 | 1,572 | 1,945 |  |
| miR-194-5p | 16,226 | 22,985 | 22,000 | 23,606 | + |
| miR-23a-3p | 10,642 | 19,591 | 23,136 | 23,473 |  |
| miR-30d-5p | 14,513 | 13,128 | 19,517 | 23,363 |  |
| miR-27a-3p/27b-3p | 12,376 | 17,883 | 13,596 | 19,461 |  |
| miR-10a-5p | 17,716 | 16,690 | 14,644 | 11,758 |  |

### Similar expression patterns among epithelial cells of different origins

Having established that a general enrichment for epithelial miRNAs occurred in the xMD-derived data, we then evaluated how similar/dissimilar our xMD-miRNA-seq derived epithelial cell miRNA expression profiles were to a colonic enteroid (colonoid) and a flow-sorted epithelial cell small RNA-seq sample. In addition, these three epithelial cell types were compared to a fibroblast, endothelial cell, lymphocyte, red blood cell and macrophage miRNA RNA-seq expression set that served as an outgroup. All of these cell types were present in the colon samples we performed xMD on and could represent potential contaminants. We selected seven known epithelial cell-enriched miRNAs and eight miRNAs that are both abundant and enriched in these non-epithelial cell types. As seen, the epithelial cell data generated from all three different cell selection methods show similar estimates for the selected miRNAs, with abundant expression of known epithelial-cell-enriched miRNAs and a paucity of non-epithelial cell-enriched miRNAs (Figure 2).

**Figure 2:**
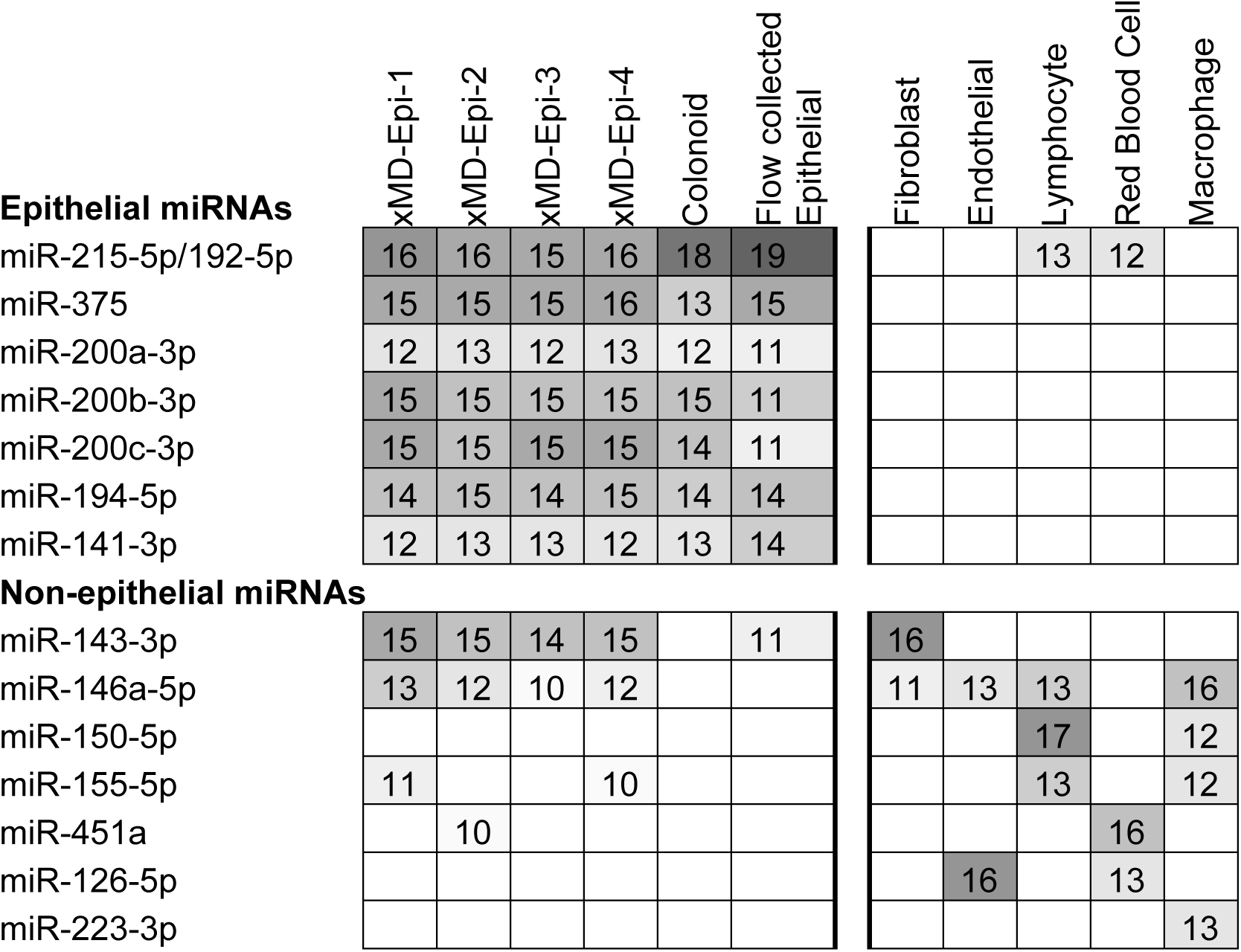
Comparison of miRNA expression patterns between epithelial cells and non-epithelial cells. Epithelial cells are enriched for 7 miRNAs that are generally not expressed in non-epithelial cells, while the xMD-miRNA-seq derived samples are generally devoid of non-epithelial specific miRNAs except for miR-143-3p. Values are log_2_ RPM rounded to a whole number. Log_2_ RPM value < 10, representing miRNAs with an RPM value < 725 were omitted for clarity. Heat map intensity corresponds to log_2_ RPM values.

### xMD-miRNA-Seq derived epithelial cells cluster with other epithelial cells

We then explored how these epithelial xMD-miRNA-seq cells would cluster among many other primary cell types, based on the small RNA-seq data. We utilized 165 primary cell small RNA-seq datasets and these four xMD-miRNA-seq samples. All samples were processed similarly in miRge and RUV normalized to correct for batch effects. Using t-SNE, we observed that cells clustered by general cell class (epithelial, mesenchymal, neural, inflammatory, red blood cell and platelet) with the xMD-miRNA-Seq derived epithelial cells clustered adjacent to other epithelial cells (Figure 3).

**Figure 3:**
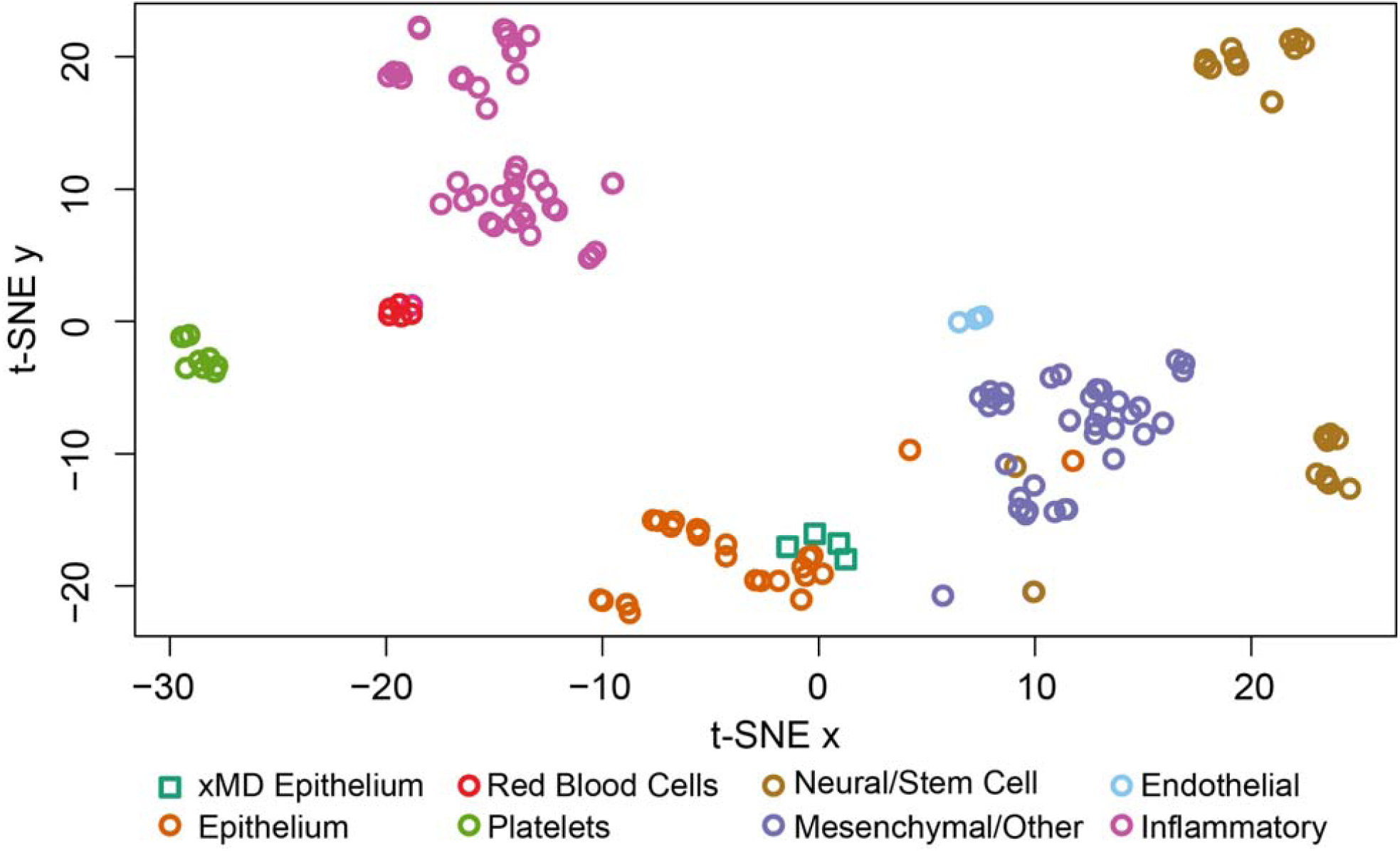
t-SNE analysis of 169 primary cells. xMD-miRNA-seq derived epithelial cells clustered adjacent to other epithelial cells and away from other cell types.

## Discussion

We introduce xMD-miRNA-seq as a method to obtain robust, cell-specific miRNA expression data from FFPE tissues. Using this method, we were able to capture a near *in vivo* signal of colonic epithelial cells from two samples. Their expression levels were consistent with miRNA expression obtained from two additional primary epithelial cell sources. Our xMD strategy successfully collected epithelial cells from the mixed colon cell population resulting in a high enrichment of epithelial cell miRNA signals with only minimal contamination from other cells of the colon.

This method has significant advantages over current options. xMD is cost-effective primarily due to extremely low upfront machine costs with mainly routine immunohistochemistry costs to isolate the cells of interest. It requires little expertise and is easily set up. Although IHC can be challenging, particularly for exotic proteins, that is not necessarily true for xMD. Well-established antibodies to perform IHC exist for most cell types, as noted at the Human Protein Atlas (proteinatlas.org).^29^ These antibodies are all that is needed to isolate specific cell types, although antibodies to more exotic proteins could further subdivide cell types (ex. FOXP3). Beyond the xMD isolation steps, the other aspects of the protocol employ routine commercially-available reagents. Thus, a high level of consistency should exist between users of the method.

While an exciting proof-of-concept has been achieved, there is room for more development and improvement across the technique. We continue to maximize the enrichment of cellular material from the FFPE slide and minimize contamination during the xMD step. Whereas six slides were used to generate the starting material for miRNA-seq, from a cost-perspective, it would be ideal to reduce that to a single slide. We also note that RNA was degraded during the IHC steps due to the introduction of RNAses in the antibody reagents and high temperature antigen retrieval (HTAR). To overcome this, we are developing methods to reduce this loss. Finally, we utilized a new NEXTflex RNA-seq library preparation method, which worked well in obtaining appropriate relative miRNA levels. However, the levels of rRNA and tRNA fragments dominated the specimens, likely due to RNA degradation and a known challenge of FFPE tissues.^30^ The percent miRNAs in any RNA-seq method should be 10% or higher, or else very deep sequencing, as performed here, will be required to obtain sufficient coverage. Additional steps to deplete rRNAs and tRNAs may be necessary to increase the miRNA yield for each sample, and thus allow for the use of this method with lower sequencing depth to reduce costs. We do note that generating 31 million+ reads/sample indicates that the amount of RNA obtained was not rate limiting, despite current inefficiencies of the method.This data highlights one of the main challenges in small RNA-seq: cross methodologic differences in sequencing libraries.^31^ The Illumina TruSeq small RNA library preparation is the most commonly used method and the source of most primary cell miRNA RNA-seq data in the literature.^3-5^ However, the method requires larger RNA inputs than are available for xMD-miRNA-Seq and has known biases, particularly in regards to adapter ligation.^32^ The NEXTflex method utilizes randomized adapters, which have reduced adapter ligation bias, and shows a different distribution of miRNA abundance with potential other biases.^33,34^ More specifically, we have found Illumina libraries tend to have a couple of dominant miRNAs (with RPM values >100,000) representing a major percentage of the reads. Conversely, in our NEXTflex data, no single miRNA was >100,000 RPM (>10% of all reads) (Table 2). There was a more even distribution of abundance across the miRNAs. Thus, a comparison across these methods may not be as robust as expected.^35^ Since this project began on the premise of finding the most accurate *in vivo* cellular miRNA expression estimate for deconvoluting tissues, this will remain a challenge if tissue data is generated in a library format that is not fully compatible with the library preparation method of xMD-miRNA-seq. In fact, while the levels of miR-192/215 were 294,000 and 184,000 in two colon samples sequenced using the Illumina system (SRR837836, SRR837842) the RPM value for miR-192/215 was lower, at 44,000-70,000 in the xMD-Epi samples.^4^ This RPM value remains too low to explain the high signal from the colon and would challenge deconvolution algorithms. Thus, having cell and tissue data matched by library preparation method, may be necessary for optimal deconvolution studies.

Using FFPE tissues for xMD-miRNA-seq has its advantages and disadvantages. Although RNAs degrade in fixed tissues over time, unlike mRNAs, miRNAs generally maintain their expression^36,37^ and correlate well with frozen tissues.^38,39^ But, miRNA degradation does appear to vary between specific miRNAs with %GC being implicated, suggesting biases in FFPE-derived data.^25^ Additionally, RNA quality is often an issue for FFPE tissue due to cross-linking. As well, devitalization times, times to fixation, or postmortem intervals (for autopsy-derived tissues), diminish the quality of the RNA before formalin fixation and storage has even begun for human tissues. These issues are independent of the cross-linking effect and must be overcome by more proactive efforts to accelerate the fixation of tissues.

Nonetheless, the ability to generate robust miRNA RNA-seq data directly from cells obtained from FFPE tissues has broad implications. Now, for the first time, we can access mature primary cell types that do not grow in culture. We can obtain the miRNA expression levels for fully mature cells that are in appropriate environments and maintaining cell-cell interactions. As well, stored FFPE tissues are abundant in pathology laboratories worldwide. Thus, one could mine this resource for just about any expression pattern of any cell type in any disease state envisioned. For example, one could isolate just CD4+ T cells from FFPE tissues of a specific tumor microenvironment to explore their miRNA expression patterns and compare these patterns to that of other T cell subsets or T cells taken from a different tissue. The possibilities to further miRNA expression analysis in a variety of disease states are enormous.

## Conclusion

This proof-of-concept study shows xMD-miRNA-seq to capture sufficient and specific quantities of a specific cell type for small RNA-seq. This method is the most cost-effective and widely applicable method yet to obtain accurate near *in vivo* miRNA expression level estimates from primary cells.

## Methods

### Expression microdissection (xMD)

Histologic sections of formalin-fixed paraffin-embedded (FFPE) sections (4 micron thick) of colon were prepared. Samples were obtained from two anonymous subjects of normal bowel obtained from surgery for a colonic mass lesion. Tissue collection was done in accordance with and approved by the Investigational Review Board of Johns Hopkins Hospital. Sections were immediately placed on DRIERITE^TM^ desiccant in a vacuum desiccator cabinet until IHC was performed. Immunohistochemistry was performed as follows: Slides were baked for 20 minutes at 60°C. Paraffin was removed from cooled slides by immersion in two changes of xylene followed by graded washes of ethanol and water to rehydrate tissues. Antigen retrieval was performed by immersing the slides in Trilogy (Sigma-Aldrich, St. Louis, MO) in a pressure cooker to 126°C and 18-23 psi. Slides were stained for cytokeratins AE1/AE3 (Diagnostic Biosystems, Pleasanton, CA) followed by incubation with a polymer HRP IgG (Leica Biosystems, Buffalo Grove, IL.) Protector RNase Inhibitor (Roche Diagnostics, Mannheim, Germany) was added to both primary and secondary antibody incubations at a concentration of 1U/μl.The antibody complex was detected with Deep Space Black (Biocare Medical. Concord California). After dehydration with xylene, slides were air dried and then stored in Drierite (W.A. Hammond Drierite Co., Xenia, OH.). xMD was performed following previously described methods.^20^ Briefly, the slides were covered with ethylene vinyl acetate (EVA) polymer film (9702, 3M) and vacuum sealed using a FoodSaver bag system. Sensepil lamp (HomeSkinovations) was used to irradiate the tissue through a blotting paper diffuser using 5 discharges at intensity level 5 (Supplementary Figure 2).

### RNA isolation

RNA was extracted from the EVA films using Qiagen miRNeasy FFPE kit. Briefly, the films were completely submerged in PKD buffer: proteinase K (15:1) and heated for 15 min at 56°C with shaking at 500 RPM. The proteinase K was quenched for 15 min at 80°C and then immediately placed on ice for 3 minutes. Following 15 minutes of centrifugation at maximum speed, DNAse booster was added (1:10) and incubated at room temperature for 15 minutes. Buffer RBC was added (2:1) and the sample passed over the Qiagen column and washed. RNA was eluted in 20 μL RNase free water. RNA was quantified using the high sensitivity RNA assay on a Qubit 3.0.

### RNA-seq for xMD derived cells

Library preparation and sequencing of the xMD samples was performed at the Lieber Institute for Brain Development. Libraries were created from 50ng of the total RNA extracted from the xMD samples using the BIOO Scientific NEXTflex library preparation kit V3 for Illumina. Small RNAs were size selected using a PAGE gel. Eighteen cycles were performed for PCR amplification. Libraries were generated using the Perkin Elmer Sciclone G3 machine using the Maestro Software version 6.0.13.152 and following the Sciclone NGS and NGSx workstation automation guide for this kit, which is downloadable from www.biooscientific.com. Libraries were sequenced with 50 base pair single-end reads using the Illumina HiSeq 3000, in a single lane with no other samples. The Illumina Real Time Analysis (RTA) module was used to perform image analysis and base calling and the BCL Converter (CASAVA v1.8.2) was used to generate the sequence reads. A sequencing depth of greater than 31 million reads for each sample was obtained, (Table 1). Adapters and the first and last 4 bases were trimmed from the FASTQ files using Cutadapt version 1.14.^40^

### Growing Enteroids

Colonic enteroids (colonoids) were grown from colonic tissue as previously described in detail.^41^ Briefly, collected colonic crypts containing stem cells were embedded in matrigel submerged in growth media containing Advanced Dulbecco’s modified Eagle medium/Ham’s F-12, 100 U penicillin/streptomycin, 10 mM HEPES, and 0.2 mM GlutaMAX, 50% v/v Wnt3a conditioned medium, 20% v/v R-spondin-1 conditioned medium, 10% v/v Noggin conditioned medium, 1x B27 supplement minus vitamin A, 1x N2 supplement, 1 mM *N*-acetylcysteine, 50 ng/mL human epidermal growth factor, 1 μg/mL [Leu-15] gastrin, 500 nM A83-01, 10 μM SB202190 [4-[4-(4-fluorophenyl)-5-pyridin-4-yl-1,3-dihydroimidazol-2-ylidene]cyclohexa-2,5-dien-1-one], 10 nM prostaglandin E2, 100 μg/mL primocin, 10 μM CHIR99021, and 10 μM Y-27632. Colonoids were passaged every 7-10 days. RNA was isolated using the miRNeasy Mini kit (Qiagen).

### Flow sorted colonic epithelial cells

Colonic epithelial cells were obtained by the flow sorting (BD FACSAria II) of EpCAM+ cells through a modification of the protocol of Dalerba.^42^ RNA was isolated using the miRNAeasy Mini kit. Small RNA-seq data from these cells were previously published in McCall et al.^4^ and are in the Sequence Read Archive as SRR5127219.

### Illumina TruSeq Small RNA-seq

Small RNA libraries of colonoids and flow sorted epithelium were prepared using the Illumina TruSeq Small RNA Library Preparation kit according to the manufacturer’s protocol, as described.^4^ Multiplexed sequencing was performed as single-read 50 bp, using rapid run mode and v2 chemistry on a HiSeq 2000 system (Illumina).

### Additional small RNA-seq data

RPM data for additional primary cells was obtained from the NCBI BioProject PRJNA358331 (SRA accession numbers SRR5127200-36 and SRR5139121) of McCall et al.^4^ This includes the flow-sorted epithelial cell described in further detail above. Other specific comparison samples used below were SRR5127200 (lymphocyte), SRR5127232 (red blood cell), SRR5127228 (endothelial), and SRR5127231 (fibroblast).

### Small RNA-seq data processing

All samples were processed using miRge^22^ or miRge 2.0^21^, an updated version of miRge. Both miRge and miRge 2.0 utilized custom expression libraries to which collapsed reads are aligned using Bowtie.^43^ To match BioProject PRJNA358331 data for the purposes of RUV normalization, we processed the four new xMD-miRNA-seq samples using the original miRge. miRNAs that were parts of repeat elements (ex. miR-5100) or coding regions of genes (ex. miR-7704), which were over-represented in this analysis, were removed. We also processed those four samples along with individual colonoid, macrophage, fibroblast, endothelial cell, red blood cell and flow-sorted epithelial cells with miRge 2.0. Using miRge 2.0, we utilized a MirGeneDB 2.0 human library containing 586 miRNA genes^23^, instead of a miRBase v21 library containing 2,588 miRNAs^24^, to focus on the most accurate list of miRNAs. Reads for each sample are reported as absolute read counts and RPM. Somewhat unique to both miRge versions, when two miRNAs are highly similar in that isomiR reads could align to either, the two (or more) miRNAs are clustered together (ex. let-7a-5p/7c-5p). Additionally, after miRge 2.0 processing, the data was hand-curated to look for anomalous miRNAs and five miRNAs (miR-328-3p, miR-34c-5p, miR-330-3p, miR-34a-5p, and miR-34c-5p) were removed from the xMD-miRNA-seq samples due to extreme skewing towards non-canonical miRNAs. As an example, in one specimen (xMD-Epi-2), miR-328-3p had only 25 canonical sequences, but had 6,495 non-canonical isomiR sequences indicating only 0.4% of reads were canonical and likely alignment of non-miR-328-3p sequences to this miRNA. A typical miRNA has >60% of reads being canonical.^4^

### Remove Unwanted Variation (RUV) normalization

We utilized the remove unwanted variation algorithm^44^ (RUVSeq R/BioC package version 1.8.0) to normalize 165 primary cell small RNA-seq runs from McCall et al.^4^ and 4 xMD-miRNA-seq samples.

### Assessing technical variation

To compare miRNA length distribution differences, the length of each canonical and non-canonical (isomiR) sequence was determined for 5 samples: xMD-Epi-2, xMD-Epi-3, a flow collected epithelial cell sample (SRR5127219), a cultured primary endothelial cell sample (SRR5127228) and a cultured primary fibroblast cell sample (SRR5127231). These values were converted to the percent of each length using a histogram function in Excel.

To assess which miRNAs varied the most between technical replicates we took the minimal and maximal RPM values for each miRNA across the two technical replicate xMD-miRNA-seq samples (1 vs. 2 and 3 vs. 4). Then we compared the fold change between these two variables among the 90 miRNAs with a minimum average value of 1,000 RPM. To calculate a coefficient of variation, the same 90 miRNAs were used. These were plotted as coefficient of variation vs. log_2_ RPM. There is again a trend for a higher coefficient of variation for the lower abundance miRNAs. However, there are no miRNAs that are particular outliers. This figure has been added to Supplementary Figure 1.

### t-Stochastic Neighbor Embedding

t-SNE was performed using the Rtsne package (version 0.11) in R on RUV-corrected cell data. For t-SNE, perplexity was set to 10.^45^

### Comparison of miRNA expression by known cell-specific patterns

Epithelial-enriched and other cell type-specific miRNAs were identified from recent publications on cell-specific miRNA expression patterns.^4,5^

### Statistical Methods

Pearson’s pairwise correlations were generated using the correlation data analysis tool in Excel on log_2_ adjusted RPM data.

## Data Availability

The four small RNA-seq datasets on the xMD-Epithelial data and the one colonoid RNA-seq dataset have been submitted to the Sequence Read Archive as BioProject PRJNA445477 samples SRR6895202-5 and SRR6898464.

## Acknowledgements

AZR, MNM, and MKH were supported by the National Institutes of Health [1R01HL137811]. MKH was supported by the American Heart Association [Grant-in-Aid 17GRNT33670405]. MNM was supported by the National Institutes of Health [R00HG006853 and UL1TR002001]. The authors thank Jennifer Foulke-Abel for her assistance in growing the colonoids.

## Authors Contributions

A.Z.R. and M.K.H. conceived of the project. C.W., A.Z.R. and M.K.H. wrote the manuscript. K.F.T., A.Z.R., C.P., O. K., C.W., A.R., C.W., J.H.S. performed experiments. M.N.M. and M.K.H. analyzed data.

## References

1. Mendell, J. T. & Olson, E. N. MicroRNAs in stress signaling and human disease. Cell 148, 1172–1187, doi:10.1016/j.cell.2012.02.005 (2012).

2. Fromm, B. et al. A Uniform System for the Annotation of Vertebrate microRNA Genes and the Evolution of the Human microRNAome. Annual review of genetics 49, 213–242, doi:10.1146/annurev-genet-120213-092023 (2015).

3. Juzenas, S. et al. A comprehensive, cell specific microRNA catalogue of human peripheral blood. Nucleic Acids Res 45, 9290–9301, doi:10.1093/nar/gkx706 (2017).

4. McCall, M. N. et al. Toward the human cellular microRNAome. Genome Res 27, 1769–1781, doi:10.1101/gr.222067.117 (2017).

5. de Rie, D. et al. An integrated expression atlas of miRNAs and their promoters in human and mouse. Nature biotechnology 35, 872–878, doi:10.1038/nbt.3947 (2017).

6. Halushka, M. K. MicroRNA-144 is unlikely to play a role in bronchiolitis obliterans syndrome. The Journal of heart and lung transplantation: the official publication of the International Society for Heart Transplantation 35, 543, doi:10.1016/j.healun.2016.01.008 (2016).

7. Kent, O. A., McCall, M. N., Cornish, T. C. & Halushka, M. K. Lessons from miR-143/145: the importance of cell-type localization of miRNAs. Nucleic Acids Res 42, 7528–7538, doi:10.1093/nar/gku461 (2014).

8. Wanjare, M., Kuo, F. & Gerecht, S. Derivation and maturation of synthetic and contractile vascular smooth muscle cells from human pluripotent stem cells. Cardiovasc Res 97, 321–330, doi:10.1093/cvr/cvs315 (2013).

9. Lopes-Ramos, C. M. et al. Regulatory network changes between cell lines and their tissues of origin. BMC Genomics 18, 723, doi:10.1186/s12864-017-4111-x (2017).

10. Kuosmanen, S. M., Kansanen, E., Sihvola, V. & Levonen, A. L. MicroRNA Profiling Reveals Distinct Profiles for Tissue-Derived and Cultured Endothelial Cells. Scientific reports 7, 10943, doi:10.1038/s41598-017-11487-4 (2017).

11. Shen-Orr, S. S. & Gaujoux, R. Computational deconvolution: extracting cell type-specific information from heterogeneous samples. Current opinion in immunology 25, 571–578, doi:10.1016/j.coi.2013.09.015 (2013).

12. McCall, M. N., Illei, P. B. & Halushka, M. K. Complex Sources of Variation in Tissue Expression Data: Analysis of the GTEx Lung Transcriptome. American journal of human genetics 99, 624–635, doi:10.1016/j.ajhg.2016.07.007 (2016).

13. Avila Cobos, F., Vandesompele, J., Mestdagh, P. & De Preter, K. Computational deconvolution of transcriptomics data from mixed cell populations. Bioinformatics, doi:10.1093/bioinformatics/bty019 (2018).

14. Newman, A. M. et al. Robust enumeration of cell subsets from tissue expression profiles. Nature methods 12, 453–457, doi:10.1038/nmeth.3337 (2015).

15. Halushka, M. K., Fromm, B., Peterson, K. J. & McCall, M. N. Big Strides in Cellular MicroRNA Expression. Trends in genetics: TIG 34, 165–167, doi:10.1016/j.tig.2017.12.015 (2018).

16. Cheng, L. et al. Laser-assisted microdissection in translational research: theory, technical considerations, and future applications. Applied immunohistochemistry & molecular morphology: AIMM 21, 31–47, doi:10.1097/PAI.0b013e31824d0519 (2013).

17. Schwarz, E. C. et al. Deep characterization of blood cell miRNomes by NGS. Cellular and molecular life sciences: CMLS 73, 3169–3181, doi:10.1007/s00018-016-2154-9 (2016).

18. Faridani, O. R. et al. Single-cell sequencing of the small-RNA transcriptome. Nature biotechnology 34, 1264–1266, doi:10.1038/nbt.3701 (2016).

19. Kumar, B. et al. Cell-type specific expression of oncogenic and tumor suppressive microRNAs in the human prostate and prostate cancer. BioRxiv **10.1101/251090**, doi:10.1101/251090 (2018).

20. Rosenberg, A. Z. et al. High-Throughput Microdissection for Next-Generation Sequencing. PLoS One 11, e0151775, doi:10.1371/journal.pone.0151775 (2016).

21. Lu, Y., Baras, A. S. & Halushka, M. K. miRge 2.0: An updated tool to comprehensively analyze microRNA sequencing data. bioRxiv **10.1101/250779**, doi:10.1101/250779 (2018).

22. Baras, A. S. et al. miRge -A Multiplexed Method of Processing Small RNA-Seq Data to Determine MicroRNA Entropy. PLoS One 10, e0143066, doi:10.1371/journal.pone.0143066 (2015).

23. Fromm, B. et al. MirGeneDB2.0: the curated microRNA Gene Database. BioRxiv, https://doi.org/10.1101/258749, xdoi:https://doi.org/10.1101/258749 (2018).

24. Kozomara, A. & Griffiths-Jones, S. miRBase: annotating high confidence microRNAs using deep sequencing data. Nucleic Acids Res 42, D68–73, doi:10.1093/nar/gkt1181 (2014).

25. Kakimoto, Y., Tanaka, M., Kamiguchi, H., Ochiai, E. & Osawa, M. MicroRNA Stability in FFPE Tissue Samples: Dependence on GC Content. PLoS One 11, e0163125, doi:10.1371/journal.pone.0163125 (2016).

26. Biton, M. et al. Epithelial microRNAs regulate gut mucosal immunity via epithelium-T cell crosstalk. Nature immunology 12, 239–246, doi:10.1038/ni.1994 (2011).

27. Chivukula, R. R. et al. An essential mesenchymal function for miR-143/145 in intestinal epithelial regeneration. Cell 157, 1104–1116, doi:10.1016/j.cell.2014.03.055 (2014).

28. Tosar, J. P., Rovira, C., Naya, H. & Cayota, A. Mining of public sequencing databases supports a non-dietary origin for putative foreign miRNAs: underestimated effects of contamination in NGS. RNA 20, 754–757, doi:10.1261/rna.044263.114 (2014).

29. Uhlen, M. et al. Proteomics. Tissue-based map of the human proteome. Science 347, 1260419, doi:10.1126/science.1260419 (2015).

30. Buitrago, D. H. et al. Small RNA sequencing for profiling microRNAs in long-term preserved formalin-fixed and paraffin-embedded non-small cell lung cancer tumor specimens. PLoS One 10, e0121521, doi:10.1371/journal.pone.0121521 (2015).

31. Dard-Dascot, C. et al. Systematic comparison of small RNA library preparation protocols for next-generation sequencing. BMC Genomics 19, 118, doi:10.1186/s12864-018-4491-6 (2018).

32. Witwer, K. W. & Halushka, M. K. Toward the promise of microRNAs -Enhancing reproducibility and rigor in microRNA research. RNA biology 13, 1103–1116, doi:10.1080/15476286.2016.1236172 (2016).

33. Baran-Gale, J. et al. Addressing Bias in Small RNA Library Preparation for Sequencing: A New Protocol Recovers MicroRNAs that Evade Capture by Current Methods. Frontiers in genetics 6, 352, doi:10.3389/fgene.2015.00352 (2015).

34. Fuchs, R. T., Sun, Z., Zhuang, F. & Robb, G. B. Bias in ligation-based small RNA sequencing library construction is determined by adaptor and RNA structure. PLoS One 10, e0126049, doi:10.1371/journal.pone.0126049 (2015).

35. Giraldez, M. D. et al. Accuracy, Reproducibility And Bias Of Next Generation Sequencing For Quantitative Small RNA Profiling: A Multiple Protocol Study Across Multiple Laboratories. bioRxiv, doi:https://doi.org/10.1101/113050 (2017).

36. Sanchez, I. et al. RNA and microRNA Stability in PAXgene-Fixed Paraffin-Embedded Tissue Blocks After Seven Years’ Storage. American journal of clinical pathology 149, 536–547, doi:10.1093/ajcp/aqy026 (2018).

37. Hall, J. S. et al. Enhanced stability of microRNA expression facilitates classification of FFPE tumour samples exhibiting near total mRNA degradation. Br J Cancer 107, 684–694, doi:10.1038/bjc.2012.294 (2012).

38. de Biase, D. et al. miRNAs expression analysis in paired fresh/frozen and dissected formalin fixed and paraffin embedded glioblastoma using real-time pCR. PLoS One 7, e35596, doi:10.1371/journal.pone.0035596 (2012).

39. Xi, Y. et al. Systematic analysis of microRNA expression of RNA extracted from fresh frozen and formalin-fixed paraffin-embedded samples. RNA 13, 1668–1674, doi:10.1261/rna.642907 (2007).

40. Martin, M. Cutadapt removes adapter sequences from high-throughput sequencing reads. EMBnet.journal 17, 10–12, doi:http://dx.doi.org/10.14806/ej.17.1.200. (2011).

41. In, J. et al. Enterohemorrhagic Escherichia coli reduce mucus and intermicrovillar bridges in human stem cell-derived colonoids. Cellular and molecular gastroenterology and hepatology 2, 48–62 e43, doi:10.1016/j.jcmgh.2015.10.001 (2016).

42. Dalerba, P. et al. Phenotypic characterization of human colorectal cancer stem cells. Proc Natl Acad Sci U S A 104, 10158–10163, doi:10.1073/pnas.0703478104 (2007).

43. Langmead, B., Trapnell, C., Pop, M. & Salzberg, S. L. Ultrafast and memory-efficient alignment of short DNA sequences to the human genome. Genome Biol 10, R25, doi:10.1186/gb-2009-10-3-r25 (2009).

44. Risso, D., Ngai, J., Speed, T. P. & Dudoit, S. Normalization of RNA-seq data using factor analysis of control genes or samples. Nature biotechnology 32, 896–902, doi:10.1038/nbt.2931 (2014).

45. van der Maaten, L. & Hinton, G. Visualizing Data using t-SNE. Mach Learn Res 9, 2579–2605 (2008).

